# Neu5Gc binding loss of subtype H7 influenza A virus facilitates adaptation to gallinaceous poultry following transmission from waterbirds but restricts spillback

**DOI:** 10.1101/2024.01.02.573990

**Authors:** Minhui Guan, Thomas J. Deliberto, Aijing Feng, Jieze Zhang, Tao Li, Shuaishuai Wang, Lei Li, Mary Lea Killian, Beatriz Praena, Emily Giri, Shelagh T Deliberto, Jun Hang, Alicia Olivier, Mia Kim Torchetti, Yizhi Jane Tao, Colin Parrish, Xiu-Feng Wan

**Author notes:** **Correspondence:** Dr. Xiu-Feng Wan by or Dr. Thomas J DeLiberto by.

## Abstract

Migratory waterfowl, gulls, and shorebirds serve as natural reservoirs for influenza A viruses, with potential spillovers to domestic poultry and humans. The intricacies of interspecies adaptation among avian species, particularly from wild birds to domestic poultry, are not fully elucidated. In this study, we investigated the molecular mechanisms underlying avian species barriers in H7 transmission, particularly the factors responsible for the disproportionate distribution of poultry infected with A/Anhui/1/2013 (AH/13)-lineage H7N9 viruses. We hypothesized that the differential expression of N-glycolylneuraminic acid (Neu5Gc) among avian species exerts selective pressure on H7 viruses, shaping their evolution and enabling them to replicate and transmit efficiently among gallinaceous poultry, particularly chickens. Our glycan microarray and biolayer interferometry experiments showed that AH/13-lineage H7N9 viruses exclusively bind to Neu5Ac, in contrast to wild waterbird H7 viruses that bind both Neu5Ac and Neu5Gc. Significantly, reverting the V179 amino acid in AH/13-lineage back to the I179, predominantly found in wild waterbirds, expanded the binding affinity of AH/13-lineage H7 viruses from exclusively Neu5Ac to both Neu5Ac and Neu5Gc. When cultivating H7 viruses in cell lines with varied Neu5Gc levels, we observed that Neu5Gc expression impairs the replication of Neu5Ac-specific H7 viruses and facilitates adaptive mutations. Conversely, Neu5Gc deficiency triggers adaptive changes in H7 viruses capable of binding to both Neu5Ac and Neu5Gc. Additionally, we assessed Neu5Gc expression in the respiratory and gastrointestinal tissues of seven avian species, including chickens, Canada geese, and various dabbling ducks. Neu5Gc was absent in chicken and Canada goose, but its expression varied in the duck species. In summary, our findings reveal the crucial role of Neu5Gc in shaping the host range and interspecies transmission of H7 viruses. This understanding of virus-host interactions is crucial for developing strategies to manage and prevent influenza virus outbreaks in diverse avian populations.

**Author Summary:** Migratory waterfowl, gulls, and shorebirds are natural reservoirs for influenza A viruses that can occasionally spill over to domestic poultry, and ultimately humans. The molecular mechanisms underlying interspecies transmission and adaptation, particularly between wild birds and domestic poultry, remain poorly understood. This study showed wild-type H7 influenza A viruses from waterbirds initially bind to glycan receptors terminated with N-Acetylneuraminic acid (Neu5Ac) or N-Glycolylneuraminic acid (Neu5Gc). However, after enzootic transmission in chickens, the viruses exclusively bind to Neu5Ac. The absence of Neu5Gc expression in gallinaceous poultry, particularly chickens, exerts selective pressure, shaping influenza virus populations, and promoting the acquisition of adaptive amino acid substitutions in the hemagglutinin protein of H7 influenza A viruses. This results in the loss of Neu5Gc binding and an increase in virus transmissibility in gallinaceous poultry, particularly chickens. Consequently, the transmission capability of these poultry-adapted H7 viruses in wild water birds decreases. Timely intervention, such as stamping out, may help reduce virus adaptation to domestic chicken populations and lower the risk of enzootic outbreaks, including those caused by influenza A viruses exhibiting high pathogenicity.

## Introduction

At least 105 wild bird species from 26 different families have been found to harbor influenza A viruses (IAVs), with waterfowl, gulls, and shorebirds being considered the primary natural reservoirs, particularly *Anseriformes* (ducks, geese, and swans) and *Charadriiformes* (gulls, terns, and waders) (1). This wide range of wild waterbirds maintains a large genetic pool of IAVs, with a total of 16 HA and 9 NA antigenic subtypes. Sporadic spillovers of avian-origin IAVs into domestic poultry are not uncommon, especially in areas with potentially inadequate biosecurity measures. Similar spillovers have also been reported in mammals (i.e., pigs, horses, and dogs) where onward transmission occurred among these new hosts (e.g., avian-like H1N1 in pigs, avian-origin H7N7 in horses, avian-origin H3N8 in horses and dogs, avian-like H3N2 in dogs). There is also potential for these viruses to spill over to humans and other non-reservoir hosts, which creates public and veterinary health burdens.

IAVs typically replicate poorly in a new host following spillover and require adaptation to overcome barriers for efficient replication and transmission. Various host factors have been associated with host adaptation of IAVs, limiting virus reservoir host range. For example, the α2-6 linkage of sialic acids and their tissue distribution limits the ability of avian IAVs to infect humans, while α-importins, DDX17 and ANP32A, or ANP32B are involved in polymerase activities that differ among IAVs isolated from avian and mammalian species (2). However, most reports have focused on virus adaptation from avian to mammalian species, or between mammalian species; however, the molecular mechanisms of interspecies adaptation among avian species are still not fully understood.

Among all HA subtypes of IAVs, H7 is commonly isolated from wild aquatic birds including dabbling ducks, diving ducks, geese, swans, and shorebirds (1). After being introduced into domestic poultry, H7 viruses can establish and lead to recurrent outbreaks where infected poultry are not depopulated promptly. The recent epizootic in China, caused by A/Anhui/1/2013-lineage H7N9 viruses (AH/13-lineage), led to at least five waves of outbreaks in humans between 2013 and 2018, resulting in 1,567 confirmed human cases, of which 615 were fatal (3), with an increase in cases from late 2016 to early 2017 (4). Notably, this virus was primarily detected in chickens, with less frequent detections in domestic duck species or other gallinaceous poultry such as quail (5–8). A laboratory experiment confirmed that AH/13-lineage H7N9 virus can efficiently spread through direct contact among chickens but not among Pekin ducks (7). These reports suggest that AH/13-lineage H7N9 viruses have undergone adaptation and acquired effective transmission ability in certain poultry species, particularly chickens, after being introduced from wild birds.

In this study, we investigated the molecular mechanisms underlying avian species barriers in H7 transmission, particularly the factors responsible for the disproportionate distribution of chickens infected with AH/13-lineage H7N9 viruses. We hypothesized that the differential expression of N-glycolylneuraminic acid (Neu5Gc) among avian species exerts selective pressure on H7 IAVs, shaping their evolution and enabling them to replicate and transmit efficiently among gallinaceous poultry, particularly chickens. We compared the glycan binding profiles of H7 IAVs, evaluated the impact of Neu5Gc expression on virus replication and evolution, and identified adaptive mutations affecting receptor binding specificity and replication ability.

## Results

### The AH/13-lineage H7N9 virus affected chickens more than other domestic avian species

To investigate the distribution of H7 IAVs in avian populations, we downloaded all available H7 strains (n = 2,651) from public databases. We sorted them by continent and functional species categories as follows: a) gallinaceous poultry such as chicken, turkey, quail, guinea fowl, and fowl; b) waterfowl such as ducks, geese, and swans, and c) all other avian species such as ostrich, ibis, parrot, and unspecified species (Supplementary Information [SI] Table S1).

Phylogenetic analyses showed that the overall H7 viruses were grouped into Eurasian and North American lineages (SI Fig. S1a). The viruses causing epizootics in domestic poultry, such as H7N1 in Italy (1999-2000) (9), H7N7 in the Netherlands (2003) (10), H7N3 in Mexico (2012-2013) (11), and AH/13-lineage H7N9 in China (2013-2017) (3), formed unique sub-lineages that were scattered across the phylogenetic tree. In contrast, viruses that caused sporadic spillovers into domestic poultry but were promptly stamped out were represented by individual branch tips mixed with clades of viral sequences recovered from various wild bird species. The majority of data across different species categories originated from Asia and North America (SI Fig. S1b).

Of the 687 AH/13-lineage H7N9 viruses from all five waves of poultry outbreaks, 613 (89.23%) were found in chickens, while only 61 (8.88%) were reported from ducks with the majority not specifying the duck species. This is consistent with surveillance data showing that over 90% of AH/13-lineage positive samples were from chickens (12, 13). In contrast, for non-AH/13-lineage H7 viruses detected in China (n=495), only 182 (36.77%) were found in chickens, compared to 256 (51.72%) detected in ducks, encompassing both wild dabbling ducks and those of unspecified species (SI Fig. S1c).

Taken together, the majority of AH/13-lineage H7N9 viruses were detected in gallinaceous poultry, particularly chickens, whereas the other H7 viruses were generally detected in a wide range of avian species, including a variety of waterfowl and aquatic species, as well as some gallinaceous poultry.

### The AH/13-lineage H7N9 virus binds exclusively to Neu5Ac, whereas other H7 viruses from wild dabbling ducks bind to both Neu5Ac and Neu5Gc

To assess receptor binding diversity, we performed glycan microarray experiments on eight AH/13-lineage H7N9 virus strains and five strains originating from wild waterbirds (Table 1). The AH/13-lineage viruses chosen for this study represent a selection from viruses responsible for the five epizootic waves. All 13 H7 viruses tested bound to α2,3-linked (SA2-3Gal) and α2,6-linked sialic acids (SA2-6Gal) but not to non-sialic acid glycans (Fig. 1). All viruses showed high affinity for SA2-3Gal but exhibited variations in their binding to SA2-6Gal. All eight AH/13-lineage viruses demonstrated a stronger binding avidity for SA2-6Gal than the other five viruses from wild waterbirds. We further identified distinct binding patterns among these H7 viruses based on the terminal sialic acid sequence N-Acetylneuraminic acid (Neu5Ac) or N-Glycolylneuraminic acid (Neu5Gc). Specifically, all eight AH/13-lineage viruses bound exclusively to glycans terminated with Neu5Ac, but not to those terminated with Neu5Gc. In contrast, all five wild waterbird viruses tested showed strong binding affinity to glycans terminated with either Neu5Ac or Neu5Gc.

**Figure 1.**
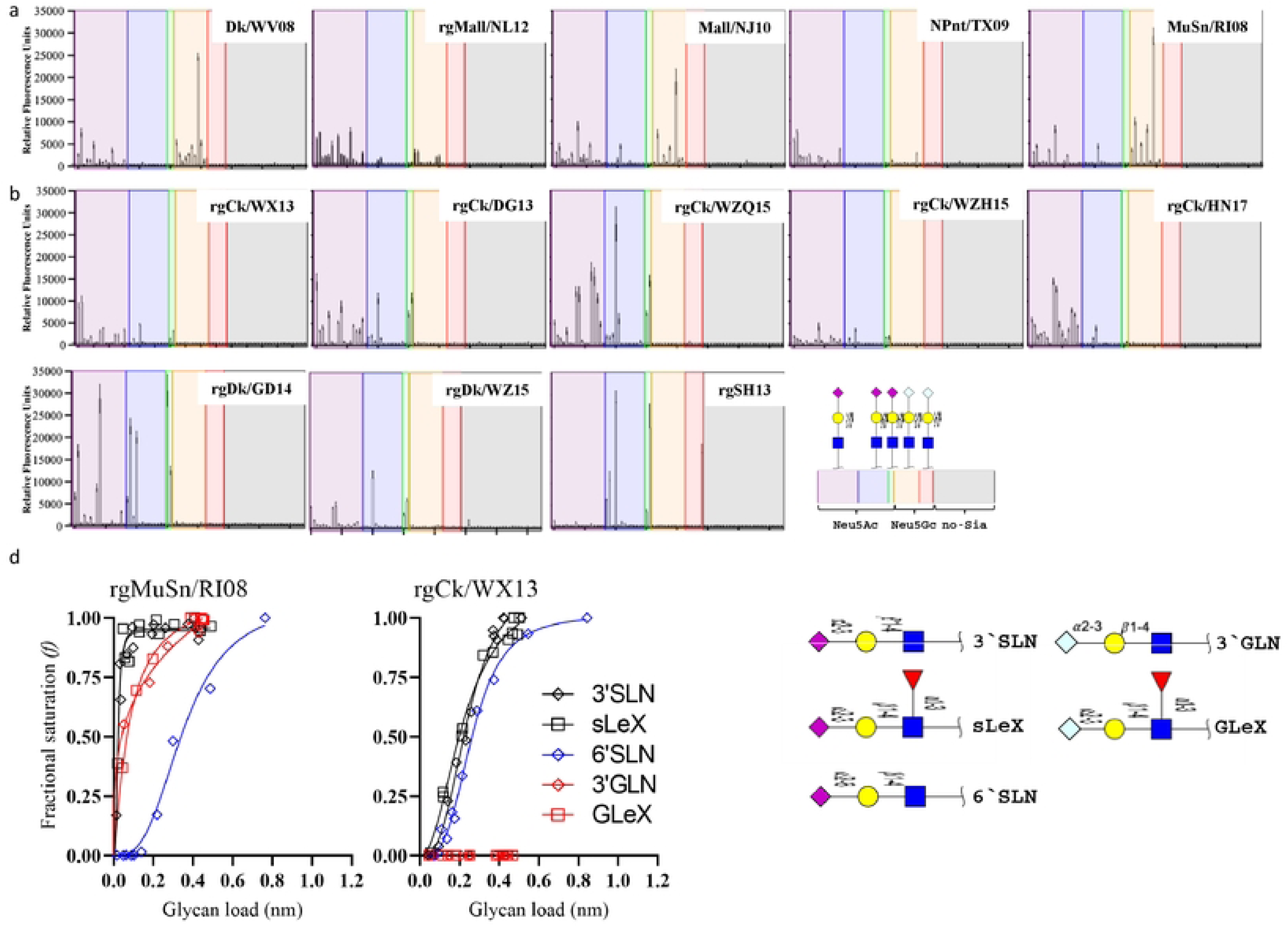
Receptor binding profile of H7 influenza A viruses. (a) N-glycan microarray binding profiles of five H7 viruses isolated from wild waterbirds in Eurasia and North America. (b) N-glycan microarray binding profiles of eight AH/13-lineage H7N9 viruses collected during the first five waves of poultry outbreaks from 2013 to 2017 in China. (c) Quantitative analyses of virus glycan binding avidity using biolayer interferometry for two representative H7 viruses, rgCk/WX13 and rgMuSn/RI08 (Table 1). We categorized 75 glycans on the microarray based on the linkage and terminal glycan sequence into α2,3- linked Neu5Ac, α2,3-linked Neu5Gc, α2,6-linked Neu5Ac, α2,6-linked Neu5Gc, and non-sialic acid glycans. The glycan sequences are detailed in SI Table S5. In the plot showing microarray data, the mean relative fluorescent units ± the standard deviations (vertical bars) are shown on the y-axis, and the x-axis represents the glycan number corresponding to the array. Biolayer interferometry analyses were performed using an Octet RED instrument (Pall FortéBio, Fremont, CA, USA) (see Materials and Methods), and binding curves were fitted using the saturation binding method in GraphPad Prism 8 (https://www.graphpad.com/scientific-software/prism/). We quantified and compared the 50% relative sugar loading concentration (RSL_0.5_) at half the fractional saturation (*f* = 0.5) of the virus against glycan analogs to determine the binding avidity. A higher RSL_0.5_ indicates a lower binding avidity.

**Table 1.** A list of subtype H7 IAVs and their corresponding abbreviations used in this study.

| Virus | Accession No for HA | Abbreviation |
| --- | --- | --- |
| **H7N9 from the domestic poultry outbreak in China (2013-2017)** |  |  |
| A/Shanghai/02/2013(H7N9) (HA,NA)× PR8<sup>a</sup> (H7N9)<sup>b</sup> | KF021597 | rgSH13 |
| A/chicken/Wuxi/0405005/2013 (H7N9) (HA)× PR8 (H7N1) | KT779570 | rgCk/WX13 |
| A/chicken/Dongguan/3418/2013(H7N9) (HA) × PR8 (H7N1) | KP413395 | rgCk/DG13 |
| A/chicken/Wenzhou/RAQL01/2015(H7N9) (HA) × PR8 (H7N1) | KU143278 | rgCk/WZQ15 |
| A/chicken/Wenzhou/HATSLG01/2015(H7N9) (HA) × PR8 (H7N1) | KU143283 | rgCk/WZH15 |
| A/chicken/Heinan/ZZ01/2017(H7N9)(HA)<sup>c</sup> × PR8 (H7N1) | MF319554 | rgCk/HN17 |
| A/Duck/Guangdong/DG527/2014(H7N9)(HA)× PR8 (H7N1) | EPI580283 | rgDk/GD14 |
| A/duck/Wenzhou/YJYF24/2015 (H7N9)(HA)× PR8 (H7N1) | ALR82230 | rgDk/WZ15 |
| **H7Nx from waterbirds** |  |  |
| A/domestic duck/West Virginia/A00140913/2008 (H7N3) | KU289983 | Dk/WV08 |
| A/mallard/Netherlands/12/2000(H7N7) (HA) × PR8-IBCDC-1 (H7N1)<sup>b</sup> | AY338460 | rgMall/NL12 |
| A/mallard/New Jersey/A00926089/2010 (H7N3) | KU290087 | Mall/NJ10 |
| A/mallard/New Jersey/A00926089/2010 (H7N3) (HA) × PR8 (H7N1) | KU290087 | rgMall/NJ10 |
| A/mute swan/Rhode Island/A00325125/2008 (H7N3) | KU290204 | MuS/RI08 |
| A/mute swan/Rhode Island/A00325125/2008 (H7N3) (HA) × PR8 (H7N1) | KU290204 | rgMuS/RI08 |
| **H7 viruses from domestic poultry (2016-2020)** |  |  |
| A/turkey/Indiana/16-001573-2/2016(H7N8) |  | Tk/IN1573-16 |
| A/duck/Alabama/17-008643-2/2017(H7N9) |  | Dk/AL8643-17 |
| A/chicken/Texas/18-007912-2/2018 (H7N1) | QKX64971 | Ck/TX7912-18 |
| A/turkey/Missouri/18-008108-11/2018 (H7N1) | QKX64983 | Tk/MO8108-18 |
| A/chicken/Missouri/18-008648/2018(H7N1) |  | Ck/MO8648-18 |
| A/turkey/California/18-031151-4/2018 (H7N3) | AYG99315 | Tk/CA1151-18 |
| A/duck/Pennsylvania/19-007197/2019(H7N3) |  | Dk/PA7197-18 |
| A/guinea fowl/Connecticut/19-009111/2019(H7N3) |  | Gf/CT9111-19 |
| A/duck/California/19-019071/2019(H7N3) |  | Dk/CA9071-19 |
| A/turkey/North Carolina/20-007949/2020(H7N3) |  | Tk/NC7949-20 |
| A/turkey/North Carolina/20-008257/2020(H7N3) |  | Tk/NC8257-20 |
| A/turkey/North Carolina/20-008425/2020(H7N3) |  | Tk/NC8425-20 |
| **Mutants** |  |  |
| A/chicken/Wuxi/0405005/2013(H7N9) (HA-A122N)× PR8 (H7N1) |  | A122N<sup>d</sup> |
| A/chicken/Wuxi/0405005/2013(H7N9) (HA-A135E)× PR8 (H7N1) |  | A135E |
| A/chicken/Wuxi/0405005/2013(H7N9) (HA-V179I)× PR8 (H7N1) |  | V179I |
| A/chicken/Wuxi/0405005/2013(H7N9) (HA-K193R)× PR8 (H7N1) |  | K193R |

<sup>a</sup>PR8, A/Puerto Rico/8/1934(H1N1); <sup>b</sup>these viruses were acquired from the BEI resources; <sup>c</sup>the PEVPRKRRTAR/GLFGA cleavage sites were mutated to PEIPKGR/GLFGA to remove multiple basic cleavage sites; <sup>d</sup> the position was based on H3 numbering.

We performed biolayer interferometry analyses for an AH/13-lineage virus, A/chicken/Wuxi/0405005/2013 (H7N9) (Ck/WX13), and a wild waterbird virus, A/mute swan/Rhode Island/A00325125/2008 (H7N3) (MuS/RI08), to further investigate the results from the glycan microarray analyses. Three testing glycan analogs, Neu5Acα2-3Galβ1-4GlcNAc (3’SLN), Neu5Acα2-3Galβ1- 4(Fucα1-3)GlcNAc (sLe^X^), and Neu5Acα2-6Galβ1-4GlcNAc (6’SLN), were terminated with Neu5Ac, whereas Neu5Gcα2-3Galβ1-4GlcNAc (3’GLN) and Neu5Gcα2-3Galβ1-4(Fucα1-3)GlcNAc (GLe^X^) were terminated with Neu5Gc. To minimize the potential impact of the neuraminidase (NA) and other gene segments, we created two reassortant viruses, rgMuS/RI08 and rgCk/WX13, each containing the HA from the corresponding parent virus aforementioned and all other segments from A/Puerto Rico/8/1934(H1N1)(PR8). Results showed that AH/13-lineage rgCk/WX13 bound exclusively to the three glycan analogs terminated with Neu5Ac but not to the others with Neu5Gc, whereas rgMuS/RI08 bound to all five analogs tested. AH/13-lineage rgCk/WX13 exhibited stronger binding avidity to 6’SLN than rgMuS/RI08. These results support the findings observed in the glycan microarray experiments (Fig. 1c).

Taken together, AH/13-lineage H7 viruses displayed a different glycan binding profile than those H7 from wild waterbirds. Specifically, AH/13-lineage viruses were found to bind to only the glycans terminated with Neu5Gc, whereas the wild waterfowl-origin H7 virus had avidity for both Neu5Ac and Neu5Gc.

### Amino acid substitution V179I expands AH/13-lineage H7 virus binding specificity from only Neu5Ac to both Neu5Ac and Neu5Gc

To identify amino acid substitutions responsible for different Neu5Gc binding patterns, we compared the HA sequences of equine H7N7, wild waterbird H7, and AH/13-lineage H7N9 viruses. Equine H7N7 were predicted to bind exclusively to Neu5Gc (14), wild waterbird H7 viruses to both Neu5Ac and Neu5Gc, and AH/13-lineage H7N9 viruses exclusively to Neu5Ac (Fig. 1). Differences were observed between AH/13- lineage H7N9 and equine H7N7 viruses, including three amino acid substitutions [i.e., I130V (H3 numbering), A135E, and K193R] in the receptor binding site (RBS) and 7 adjacent to the RBS (i.e., A122N, S128T, A160V, R172K, K173R, S174E, and V179I) (Fig. 2a).

**Figure 2.**
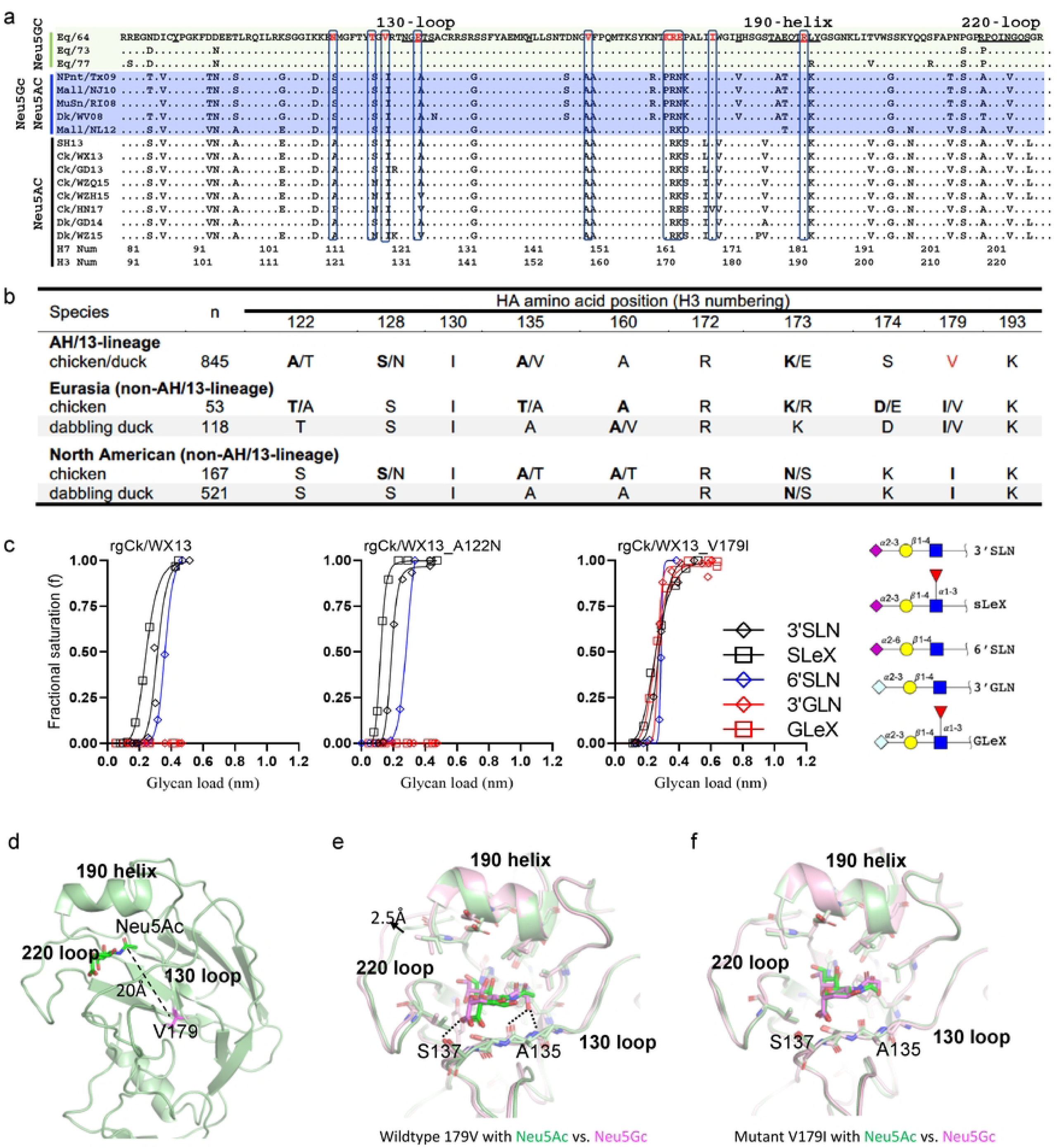
Multiple individual amino acid substitutions facilitate acquisition of virus binding avidity to Neu5Gc for H7 IAVs. (a) Sequence alignment of the receptor binding site (RBS) of H7 IAVs, including three groups of H7 viruses with distinct binding patterns to glycans terminated with Neu5Ac and Neu5Gc: equine H7N7 viruses bound exclusively to Neu5Gc, wild waterbird viruses bound to both Neu5Ac and Neu5Gc, and AH/13-lineage H7N9 viruses bound exclusively to Neu5Ac. (b) Amino acid diversity at the residues close to or within the hemagglutinin RBS of AH/13-lineage H7N9 viruses isolated from domestic poultry in China, as well as H7 viruses from dabbling ducks in Eurasian and North American (see additional details in SI Table 2). (c) Quantitative analyses of virus glycan binding avidity using biolayer interferometry for Ck/WX13, Ck/WX13-A121N, and Ck/WX13-V179I. Please refer to the legend of Figure 1 and Online Methods for the details of Biolayer interferometry analyses. (d) The crystal structure of the HA protein from A/Anhui/1/2013 (H7N9), which is identical to that of Ck/WX13. (e) Structural model of wild-type H7 in complex with Neu5Ac (green) vs. Neu5Gc (magenta). HA residues less than 3 Å away from the modeled receptor are shown in sticks. (f) Structural model of the V179I H7 mutant in complex with Neu5Ac (green) vs. Neu5Gc (magenta).

**Table 2.**
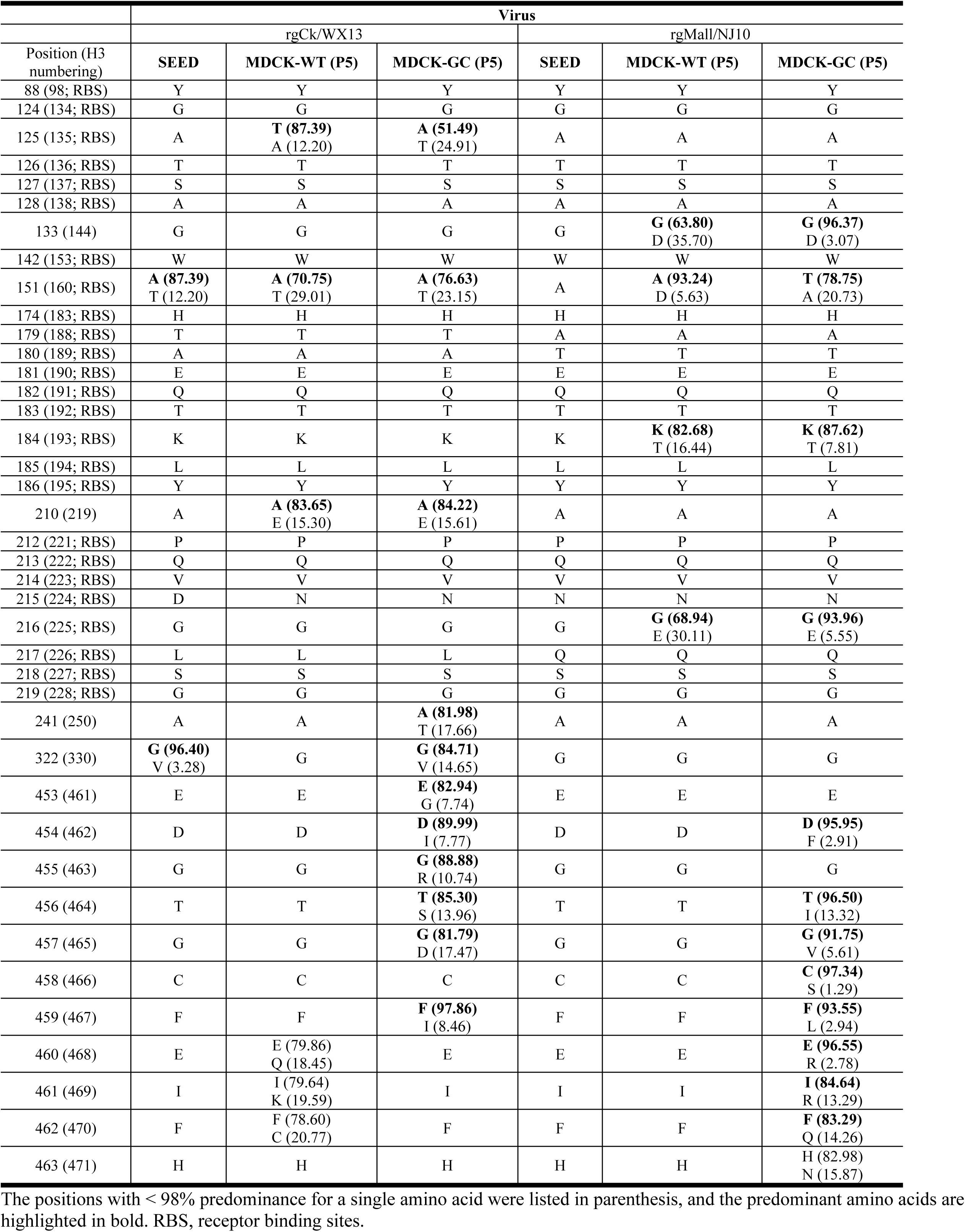
Amino acid polymorphisms in the HA protein for H7 viruses passaged in MDCK-wt and MDCK-Gc cells.

We further compared the HA sequences between AH/13-lineage H7N9 viruses and their precursor Eurasian H7 viruses isolated from wild dabbling ducks (15), specifically at the residues mentioned earlier. Amino acid substitutions were detected in the majority of AH/13-lineage isolates at residue 122 and 179 (Fig. 2b). It is noteworthy that amino acid polymorphisms were observed in all ten residues of AH/13-lineage H7N9 viruses, although to a lesser extent in wild waterbirds from both Eurasia and North America (SI Table S2). Of note, a similar V179I substitution was also observed between poultry adapted H7N7 viruses in the Netherlands (2003) and their precursor virus in wild waterbirds.

To determine the residue responsible for the loss of Neu5Gc binding ability in AH/13-lineage viruses, we replaced HA V179 (found in AH/13-lineage viruses) with I179 (present in waterfowl/equine H7-like viruses) and then created a reassortant virus rgCk/WX13-V179I. We generated three additional mutants: rgCk/WX13-A122N, rgCk/WX13-A135E, and rgCk/WX13-K193R, by substituting HA positions A122, A135, and K193 in Ck/WX13 with N122, E135, and R139, respectively. Residue 122 was included due to the significant amino acid polymorphisms (including N122) observed at this HA position in AH/13-lineage viruses. The HA mutations A135E and K193R were reported to enhance the binding of H7 virus to Neu5Gc (16). Biolayer interferometry analysis revealed that the V179I substitution conferred binding avidity of the AH/13-lineage Ck/WX13 virus to Neu5Gc while maintaining binding to Neu5Ac (Fig. 2c and SI Fig. S2). A135E and K193R substitutions, as previously reported, enhanced the binding of AH/13-lineage Ck/WX13 virus to Neu5Gc (16), whereas A122N substitution did not significantly affect the virus’ glycan binding preference to Neu5Gc or Neu5Ac.

We further performed structural modeling to understand how the amino acid substitution HA I179V enables virus binding from both Neu5Ac and Neu5Gc to Neu5Ac alone. Close inspection of the HA structure shows that V179 (found in poultry-adapted AH/13-lineage viruses) is located at the hydrophobic core of the molecule approximately 20 Å away from the RBS (Fig. 2d). Because this residue is tightly packed against several hydrophobic residues, we hypothesize that V179I, which replaces a poultry-adapted residue with residues characteristic of wild birds, would lead to conformational changes that can propagate to the 130- loop and other structural elements around the RBS and consequently broaden HA binding ability from Neu5Ac to both Neu5Ac and Neu5Gc.

Our modeling results indicate that, in the AH/13-lineage H7 (with HA-V179), Neu5Gc occupies a somewhat different position compared to Neu5Ac. Neu5Gc is shifted more towards the 220-loop in the RBS (Fig. 2e). Several close contacts (<3 Å) are observed between Neu5Gc and the RBS, including two between the extra hydroxyl group in Neu5Gc with A135 and one between the carboxyl group at the C2 position with S137 (Fig. 2e). Also, the 220-loop in the Neu5Gc structure is pushed outward by ∼2.5 Å, which is presumably necessary to accommodate Neu5Gc in the RBS. These close contacts and large structural rearrangement of the RBS needed to accommodate Neu5Gc suggest unfavorable binding. Interestingly, the modeled structures of the HA-I179, which are found in waterfowl/equine H7-like viruses, showed very similar binding modes for both Neu5Gc and Neu5Ac (Fig. 2f). In the HA-I179, there is no close contact with Neu5Gc, and the 220-loop in the Neu5Gc complex assumes a nearly identical conformation as in the Neu5Ac complex. This observation explained how the HA-I179 allows the binding to both Neu5Gc and Neu5Ac without any significant structural rearrangement.

Taken together, these results suggest that amino acid substitution V179I, which replaces a poultry-adapted residue with residues characteristic of wild birds, may facilitate the acquisition of binding avidity of AH/13- lineage H7N9 IAVs to Neu5Gc while still maintaining binding to Neu5Ac.

### Neu5Gc expression affects H7 virus replication and facilitates acquisition of adaptive mutations in the HA of H7 IAVs

We hypothesized that the expression of Neu5Gc hinders replication in H7 viruses that exclusively bind to Neu5Ac but do not affect those that bind to both Neu5Ac and Neu5Gc. To test this, we compared the growth kinetics of two mutants (rgCk/WX13-V179I and rgCk/WX13-A122N) and their parent virus, rgCk/WX13, on MDCK-Gc, a cell line expressing Neu5Gc (17), and MDCK-wt, the wild-type cell line without Neu5Gc expression. We also included A/mallard/New Jersey/A00926089/2010 (H7N3) (HA)×PR8 (H7N1) (rgMall/NJ10), with the HA of an H7 virus from a mallard (*Anas platyrhynchos*), and A/chicken/Heinan/ZZ01/2017(H7N9)(HA)×PR8 (H7N1) (rgCk/HN17), with the HA of another AH/13- lineage H7N9 virus from chicken (Table 1). All six viruses tested had identical gene segments except for the HA to minimize the impact of NA and other gene segments (see details in Online Methods). Notably, rgCk/WX13-V179I and rgMall/NJ10 bound to both Neu5Ac and Neu5Gc, whereas rgCk/HN17, rgCk/WX13, and rgCk/WX13-A122N bound exclusively to Neu5Ac.

Results indicated that both rgCk/HN17 and rgCk/WX13 viruses showed more efficient replication in MDCK-wt cells than in MDCK-Gc cells (p=0.0062 and 0.0235, respectively) (Fig. 3a). As expected, rgCk/WX13-A122N displayed similar growth kinetics as its parent virus rgCk/WX13. In contrast, the mutant rgCk/WX13-V179I showed no significant difference in growth kinetics in either cell line, similar to rgMall/NJ10, but different from their respective wild-type parent virus, rgCk/WX13.

**Figure 3.**
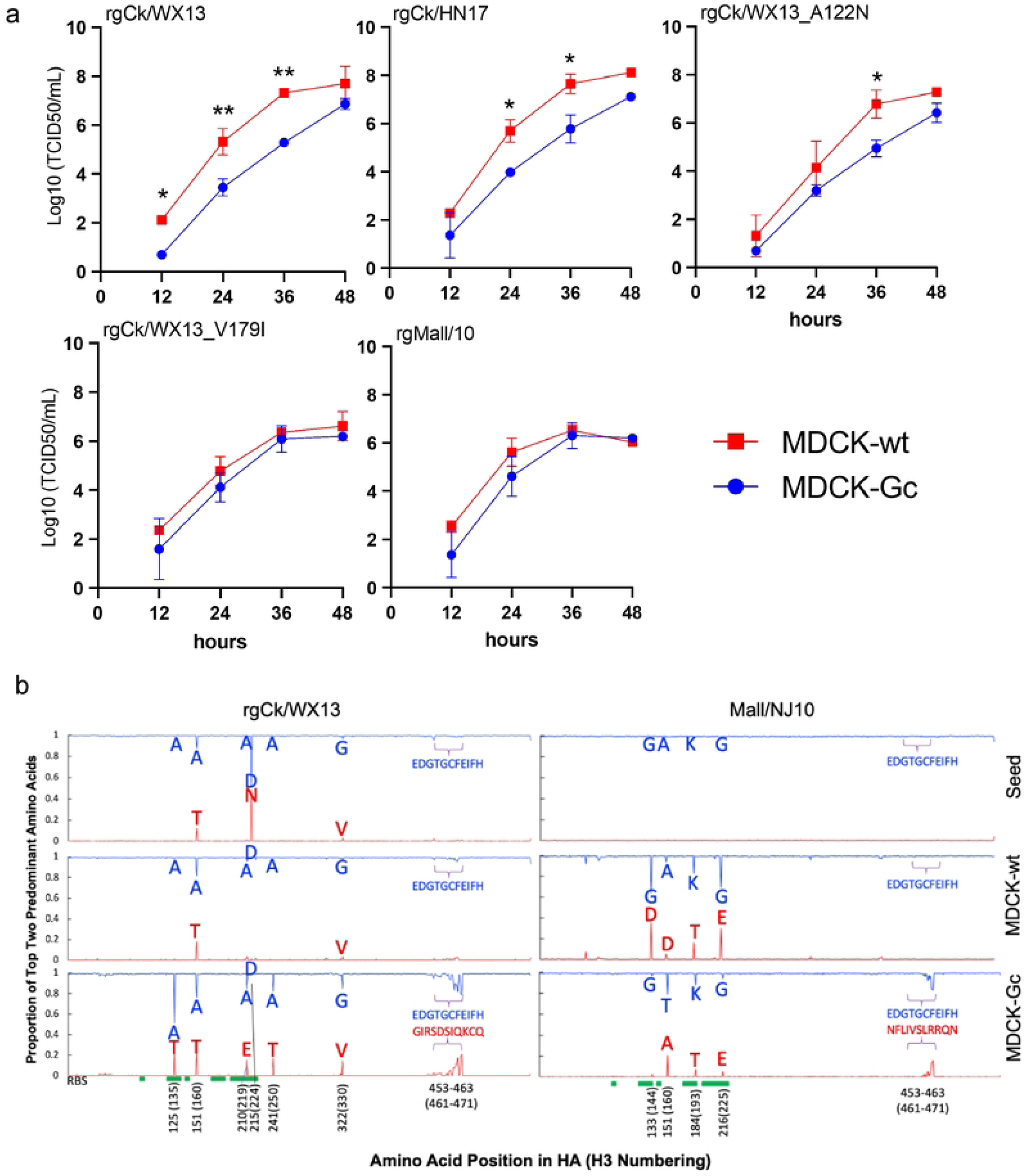
Neu5Gc affects virus replication of AH/13-lineage H7N9 viruses and drives adaptive mutations in the HA protein. a) Growth kinetics of H7 influenza A viruses and mutants in MDCK-wt and MDCK-Gc cells. All viruses had HA genes from H7 viruses and the other seven from PR8, and three mutants were generated using the HA gene of Ck/WX13 as a template. Supernatants were collected at 12-, 24-, 48-, and 72-hours post-infection (hpi) and titrated by TCID50 in MDCK CCL-34 cells. Two-way repeated measures ANOVA were used to compare time-course growth data of H7 viruses among different cells. Statistical comparisons were shown as follows: not significantly different as n.s. (P > 0.05); P ≤ 0.05 as *; P < 0.01 as **; P < 0.001 as ***; and P < 0.0001 as ****. (b) HA amino acid polymorphisms detected in the seed viruses and the viruses from the 5th passage in MDCK-wt and MDCK-Gc cells. The viruses rgCk/WX13 and rgMall/NJ10, which have an HA gene from rgCk/WX13 and rgMall/NJ10, respectively, and other seven genes from PR8 were passaged five times in MDCK-wt and MDCK-Gc cells. The two most abundant nonsynonymous mutations in the HA protein were plotted to visualize adaptive amino acid substitutions caused by cell passages. The location of the RBS in the HA protein was marked green.

To investigate whether Neu5Gc expression shapes the evolution of H7 IAVs, we passaged the rgCk/WX13 and rgMall/NJ10 in MDCK-wt or MDCK-Gc cells for five passages. For each seed, we compared their amino acid polymorphisms in HA with those in the associated fifth passage from both cells. Compared to the seed, rgCk/WX13 gained polymorphisms in MDCK-Gc at residues 135 (A to A/T), 160 (changes in A/T ratio), 219 (A to A/E), 224 (D/N to D), and 250 (A to A/T), whereas rgMall/NJ10 acquired polymorphisms in MDCK-wt at residues 144 (G to G/D), 193 (K to K/T), and 225 (G to G/E) (Table 2). Interestingly, in MDCK-Gc (but not MDCK-wt), both viruses acquired adaptive substitutions at residues 461-471 of the fusion domain, although the changes in amino acids were not identical. Overall, rgCk/WX13 exhibited a higher number of polymorphisms in MDCK-Gc compared to MDCK-wt, whereas rgMall/NJ10 had an increase in polymorphisms in MDCK-wt compared to MDCK-Gc (Fig. 3b). These findings are consistent with a prior study that demonstrated human seasonal H1N1 and H3N2 IAVs, which bind exclusively to Neu5Ac, developed adaptive HA mutations when passaged in MDCK cells expressing Neu5Gc (17). Conversely, an enzootic canine H3N2 IAV, which binds to both Neu5Gc and Neu5Ac, did not exhibit significant HA mutations under the same conditions.

Taken together, the expression of Neu5Gc hinders replication of H7 viruses that exclusively bind to Neu5Ac but does not affect those that bind to both Neu5Ac and Neu5Gc, supporting our hypothesis. In addition, Neu5Gc expression creates a selective pressure that facilitates the acquisition of adaptive substitutions in H7 IAVs, particularly those residues within or near the HA RBS.

### Distribution of Neu5Gc in the respiratory and gastrointestinal tract tissues of chicken, wild Canada goose, and selected wild dabbling duck species

To investigate Neu5Gc expression patterns in chicken, wild Canada goose, and selected wild dabbling duck species, we conducted immunofluorescence (IF) staining using a Neu5Gc-specific antibody on formalin-fixed tissues from selected avian species. We examined the trachea, small intestine (duodenum/jejunum), and large intestine (colon and cloaca) of chicken, wild Canada goose (*Branta canadensis*), and five commonly surveyed wild dabbling ducks in North American IAV surveillance: mallard, gadwall (*Mareca strepera*), green-winged teal (*Anas carolinensis*), northern shoveler (*Spatula clypeata*), and wood duck (*Aix sponsa*). Results showed distinct Neu5Gc expression patterns between the species tested (Figure 4). Neu5Gc expression was detected in mallard, green-winged teal, northern shoveler, and wood duck. In contrast, domestic chicken, Canada goose, and gadwall displayed no positive staining (Figure 4a). Among the species with positive immunostaining, Neu5Gc expression was detected in the ciliated epithelial cells of the trachea and crypt cells within the duodenum/jejunum, colon, and cloaca (Table 3). Notably, the northern shoveler exhibited significant Neu5Gc expression in the trachea, a pattern not observed in the other tested species.

**Figure 4.**
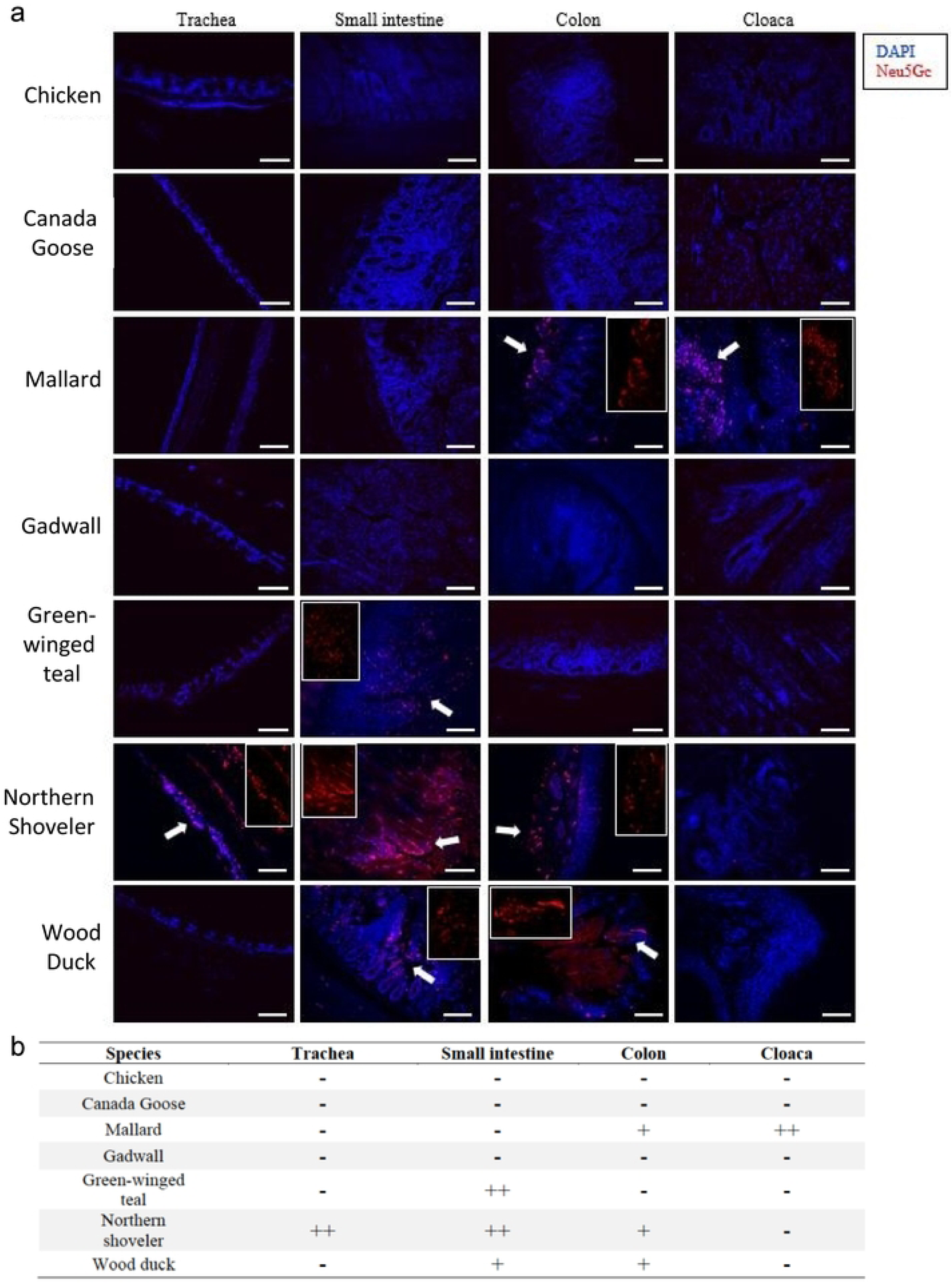
Distribution Neu5Gc glycan in the tissues of chicken, wild dabbling ducks, and Canada goose (*Branta canadensis*). Trachea, small intestine (duodenum/jejunum), colon and cloaca of chicken (*Gallus gallus*), Canada goose, mallard (*Anas platyrhynchos*), gadwall (*Mareca strepera*), green-winged teal (*Anas carolinensis*), northern shoveler (*Spatula clypeata*), and wood duck (*Aix sponsa*). Glycan terminated with Neu5Gc (red) was detected by immunofluorescence assay with anti-Neu5Gc polyclonal antibody. Nuclei were stained with DAPI (blue). The white arrows indicated positive staining of Neu5Gc, the areas of which have been enlarged at the side of each image. The scale bar at the bottom of each image was 100 µm. b) The abundance of Neu5Gc expression. We categorized the glycan receptor abundance: none or limited staining (-) without stained cells, moderate and sporadic staining (+) with <30% stained cells, and strong staining (++) with ≥ 30% of the stained cells.

In conclusion, Neu5Gc expression was absent in tissues tested from chicken, wild Canada geese, and gadwall, and was variable across the other wild duck species tested. As previously reported, all species expressed Neu5Ac (18, 19).

### A model proposed for H7 IAV adaptation and subsequent transmission upon spillover from wild waterbirds to gallinaceous poultry

This study demonstrates that AH/13-lineage H7N9 viruses in gallinaceous poultry, particularly chickens, bind exclusively to Neu5Ac, whereas H7 IAVs enzootic in wild waterbirds show binding affinities to both Neu5Gc and Neu5Ac. Strikingly, like the AH/13-lineage H7N9 viruses, all three H7 viruses responsible for recent epizootics in gallinaceous poultry also bind solely to Neu5Ac but not to Neu5Gc (16, 20, 21). These three outbreaks are: H7N1 in the chicken population of Italy (1999–2000), H7N7 in the chicken population of the Netherlands (2003), and H7N3 in the chicken population of Mexico (2012-2013). By combining the findings on virus receptor binding specificity and host Neu5Gc expression, we propose a model for H7 IAV spillover from wild waterbirds to gallinaceous poultry (Fig. 5) and the potential evolutionary impact on subsequent transmission. An H7 IAV, with binding ability to both Neu5Gc and Neu5Ac, can be transmitted efficiently among waterbird species possessing both receptors. This virus may spillover into gallinaceous poultry. Once spillover occurs, a virus may acquire specific adaptive mutations in the HA protein and then lose Neu5Gc binding specificity (with the exclusive Neu5Ac binding ability). Consequently, the transmission capability of these poultry-adapted viruses increases for gallinaceous poultry but decreases for wild waterbirds that possess both receptors. Further investigation is warranted to determine whether loss of Neu5Gc binding specificity can also occur for other subtypes of IAVs, including H5, and to understand whether Neu5Gc expression variations can restrict the transmission and adaption of IAVs among waterbirds, e.g. between those species with Neu5Gc expression and those without.

**Figure 5.**
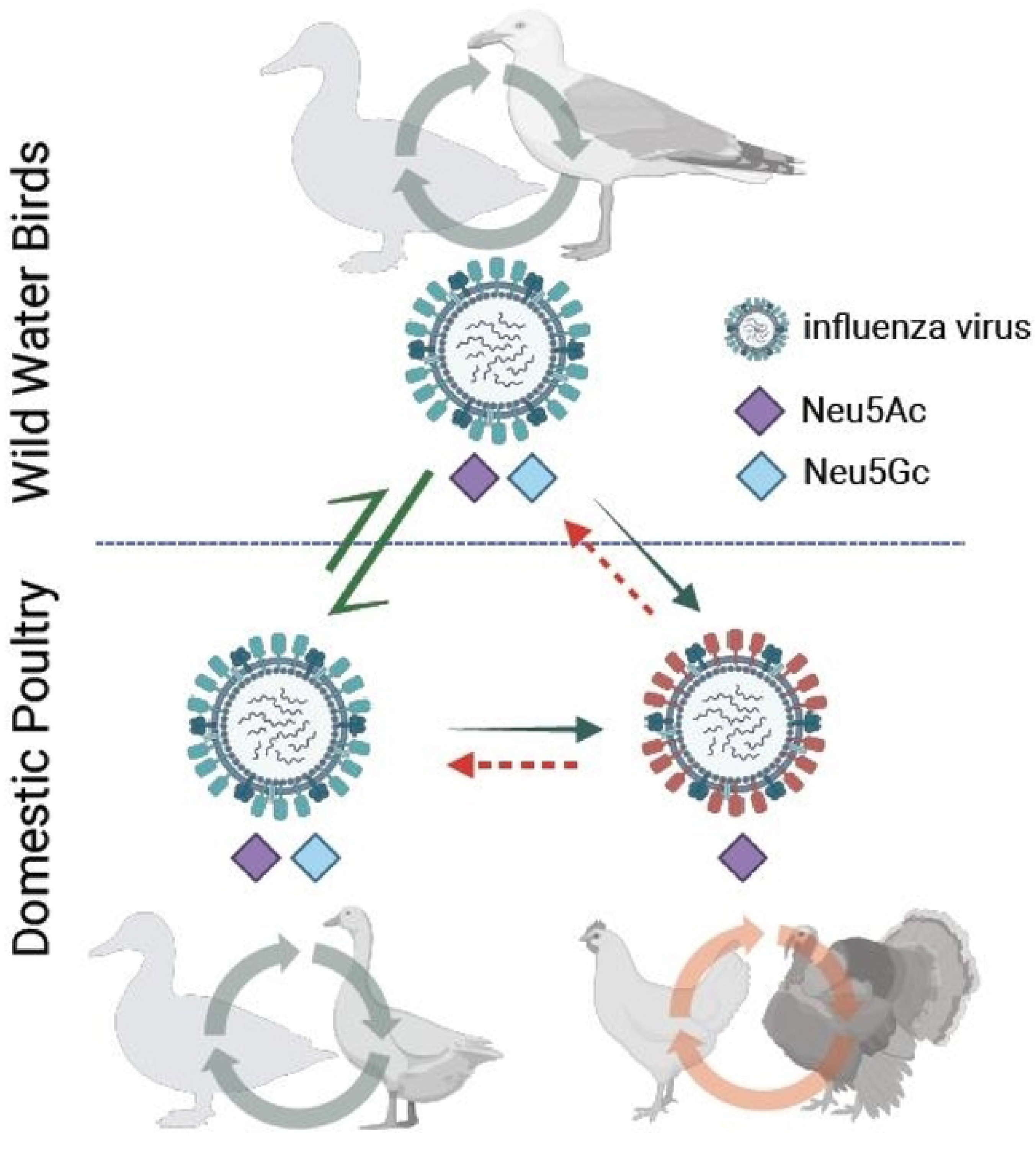
A transmission model illustrating the mechanisms of H7 IAV transmission and evolution between wild waterbirds and domestic poultry. An H7 IAV capable of binding to sialic acid receptors containing either Neu5Gc or Neu5Ac can be transmitted among wild waterbirds possessing these receptors, and can also transit between wild and domestic waterbirds expressed with the same receptors. This virus may then spill over into domestic poultry species (or another wild bird species) that expresses only Neu5Gc. Subsequent to this, the virus could acquire adaptive amino acid substitutions in the HA protein, leading it to lose its Neu5Gc binding ability and exclusively bind to Neu5Ac. Consequently, the transmission capability of these adapted viruses in waterbirds decreases.

## Discussion

A wide variety of IAVs circulate among wild waterbirds, including migratory waterfowl, gulls, and shorebirds, and occasionally transmit to domestic poultry. However, these waterbird-origin viruses typically have poor replication efficiency in gallinaceous poultry. While all avian species share the SA2- 3Gal receptor, this alone cannot explain the species barriers between waterbirds and gallinaceous poultry based on receptor binding. Among all IAVs, H7 viruses are one of the most commonly identified subtypes and often isolated from migratory waterfowl, in particular dabbling ducks, diving ducks, shorebirds, geese, and swans (SI Fig. S1) (1). This study demonstrates that wild-type H7 viruses tested from wild waterbirds have glycan binding preferences for both Neu5Gc and Neu5Ac, while those that have undergone sustained transmission in gallinaceous poultry, particularly chickens, have lost the ability to bind Neu5Gc (Fig. 1). The absence of Neu5Gc expression in chickens exerts a selective pressure on waterbird-origin H7 viruses, facilitating the acquisition of adaptive HA substitutions (20), such as I179V. Such adaptive substitutions result in the loss of Neu5Gc binding, enhance virus replication in Neu5Gc-free cells (Fig. 3), may have increased virus transmissibility in chickens (7), and consequently have led to the low detection rate of AH/13-lineage virus in domestic ducks during surveillance (12, 13). It would be useful to ascertain the point at which substitutions leading to the loss of Neu5Gc binding ability occur. This knowledge will enable timely interventions, such as depopulation, rather than resorting to controlled marketing, using this as an indicator (as illustrated in Fig. 5).

IAVs have large variations in binding avidities to Neu5Gc and Neu5Ac (22, 23), and most duck and gull origin IAVs from different subtypes including H1–H6, H9, and H11–H14 exhibited binding preference both to Neu5Gc and Neu5Ac but with stronger binding avidity to Neu5Gc (24). Thus, further studies are required to fully comprehend how Neu5Gc expression influences viral evolution and transmission of other subtypes of IAVs, particularly H5, among avian species, which could inform the development of more effective prevention and control measures against the spread of IAVs.

Infections caused by IAVs of subtypes H5 and H7 are listed as notifiable diseases, when detected in domestic poultry, by the World Organization for Animal Health due to the potential for these viruses to exhibit high-pathogenicity among domestic poultry (25). Although national programs are in place to control avian influenza in poultry worldwide, measures such as active and passive surveillance of poultry and wild birds, poultry vaccination, and stamping out of positive cases among domestic flocks, have not always achieved eradication among domestic birds (25). In the United States, stamping out has been the primary method for controlling avian influenza viruses among domestic birds, and this method has successfully controlled all introductions of H7 IAVs into poultry during the past 10 years (26). We explored genomic polymorphisms of H7 isolates from eight spillover cases in US poultry (n=18) and wild dabbling ducks (n=85) (SI Table S3), and the results indicated that H7 viruses detected in US poultry exhibited more amino acid polymorphisms compared to those found in wild dabbling ducks (SI Fig. S3). However, it is important to note that only a small subset of these polymorphisms led to substitutions at the amino acid level (SI Table S4). Notably, two out of 18 isolates exhibited the key adaptive substitution V179I; and yet, all of the strains maintained their ability to bind to both Neu5Gc and Neu5Ac, suggesting that they had not fully adapted to the domestic poultry host (SI Fig. S3c). These results imply that the prompt implementation of stamping out policies has likely been effective in preventing the emergence of poultry-adapted H7 strains. Thus, strict implementation of early detection and control policies may be crucial in minimizing the possibility of viral adaptation in poultry. That is, by promptly detecting and eliminating low pathogenicity avian influenza before they have the chance to adapt to domestic poultry, the risk of enzootic outbreaks, including potential highly pathogenic avian influenza outbreaks, can be significantly reduced.

In addition to avian species, H7s have sporadically caused infections in other mammals such as horses, harbor seals (27–29), and swine (29); some have been reported in humans as well (27, 30–38). Several previous reports support that Neu5Gc binding affects virus replication in mammals with Neu5Gc expression, although the role of Neu5Gc expression was not fully defined. For example, virus binding to Neu5Gc is a feature of IAVs that replicate in horses (39), and both an avian-origin H3N2 and an equine-origin H3N8 IAV acquired W222L, which leads to an increase in the binding affinity to Neu5Gc and enhances virus infection in canines (40, 41). On the other hand, loss of Neu5Gc binding could increase the viral infectivity to humans (42), and this could have enhanced the spillover of the AH/13-lineage H7N9 and the Netherland H7N7 (2003) viruses to humans (10).

The enzyme CMAH (cytidine monophospho-N-acetylneuraminic acid hydroxylase) converts Neu5Ac to Neu5Gc. Some mammals such as horses, dogs, and pigs appear to maintain CMAH function and predominantly express Neu5Gc in various tissues (43–46) (including the respiratory tract epithelium)(39), whereas humans and some mammals (such as seals) have lost CMAH function and do not express Neu5Gc (45, 47, 48). Genomic analyses have shown that all birds lack CMAH homologs (48), and Neu5Gc expression has not been detected in the muscle tissues in chicken, emu, and parrots; however, CMAH homologs have been detected in the liver or eggs of some avian species, possibly acquired from the diet or via an alternative pathway (49). Through an immunofluorescence assay, we demonstrated that Neu5Gc expression was widely present in the gastrointestinal tissues, such as cloaca and colon, of selected wild dabbling ducks, including mallard, green-winged teal, northern shoveler, and wood ducks, but not those of chickens (Fig. 6), which is consistent with a prior report that Neu5Gc was expressed in the intestines of Pekin ducks (*Anas platyrhynchos domesticus*) and mallards, mainly on the crypt epithelial cells of the colon (22). In addition to the gastrointestinal tracts, Neu5Gc expression was also observed in the respiratory tract tissues of the green-winged teal. Interestingly, neither the gastrointestinal nor the respiratory tracts of the gadwall or Canada goose showed Neu5Gc expression. Future research is needed to investigate Neu5Gc expression in other avian species, both wild and domestic, not included in this study, such as diving ducks, gulls and terns, snow geese, swans, sea ducks, shorebirds, and others that might serve as natural reservoirs for avian influenza viruses. Additional investigation is warranted to determine if Neu5Gc expression influences IAV transmission and adaptation between waterbird species that express Neu5Gc and those that do not.

## Materials and Methods

### Data

As of March 1, 2023, a total of 2,651 HA genomic sequences of subtype H7 avian IAVs were obtained from the Influenza Research Database (http://www.fludb.org). The sequences were sorted by location on five continents (Africa, Asia, Europe, North America, Oceania, and South America), as well as by species category. The species were categorized as gallinaceous poultry (chicken and others), waterfowl (duck and others), and other avian species (SI Table S1).

### Viruses and virus propagation

The study utilized 26 strains of IAVs, including eight genetic reassortants containing the HA gene of AH/13-lineage H7N9 viruses, and 18 wild-type or genetic reassortant viruses associated with contemporary H7 viruses from wild birds or domestic poultry from both North America and Eurasia (Table 1). All viruses were propagated in 9-day-old specific pathogen-free embryonated eggs at 37 °C for 72 hours. The resulting allantoic fluids were collected and used in growth kinetics and virus purification, or stored at -80°C until needed for analysis.

### Cells

The MDCK cells (CCL-34) were obtained from American Type Culture Collection (ATCC). The wild type MDCK NBL-2 cells (MDCK-wt) and the MDCK cells expressing Cytidine monophospho-N-acetylneuraminic acid hydroxylase (CMAH) (MDCK-Gc) were adapted from another study (17). Limited Neu5Gc (<1%) was detected in MDCK-wt whereas approximately 40% of their total Sia in MDCK-Gc as Neu5Gc (17). The cells were maintained in Dulbecco’s modified Eagle’s medium (DMEM, Gibco, New York, USA) supplemented with 10% fetal bovine serum (FBS; Atlanta Biologicals, Lawrenceville, GA, USA) at 37°C under 5% CO_2_.

### Nucleotide extraction, PCR, qRT-PCR, and genomic sequencing

Viral RNA was extracted from the allantoic fluid of embryonated chicken eggs or cell culture supernatants by using the GeneJET Viral DNA/RNA purification kit (Thermo Fisher Scientific, Waltham, MA). The RNA was subjected to cDNA synthesis using SuperScriptTM III Reverse Transcriptase (Thermo Fisher Scientific, Waltham, MA) according to the manufacturer’s instructions. PCR products of the full-length HA were generated using IAV-specific primers (50). The plasmids were then extracted with GeneJET Plasmid Miniprep Kit (Thermo Scientific, Rockford, IL). The PCR products and plasmid insertions were confirmed without unexpected mutations by using Sanger sequencing.

### Gene synthesis, molecular cloning, and reverse genetics

The HA genes of AH/13-lineage H7N9 viruses (as listed in Table 1) were synthesized and cloned into the pHW2000 vector by Gene Universal Inc. (Newark, DE). Meanwhile, the HA genes of A/mallard/New Jersey/A00926089/2010 (H7N3), A/domestic duck/West Virginia/A00140913/2008 (H7N3), and A/mute swan/Rhode Island/A00325125/2008 (H7N3) were cloned into the pHW2000 vector using a universal primer described elsewhere (50). To generate the reassortant viruses, a HA gene from a H7 virus and seven other gene segments from A/Puerto Rico/8/1934 (H1N1) (PR8) were included using reverse genetics (51) (see Table 1 for details). The nucleotide sequences of the HA gene in each rescued virus were confirmed without unexpected mutations by Sanger sequencing.

### Site-directed mutagenesis

To identify the specific amino acid substitution responsible for the acquisition of viral binding avidity to a sialic acid glycan terminated with Neu5Gc, we utilized the HA gene of Ck/WX13 as a template to create a set of mutants by site-directed mutagenesis. These mutants included amino acid substitution A122N (H3 numbering), A135E, V179I, or K193R in HA protein. To generate a specific mutation in the HA gene of Ck/WX13, we used the Phusion™ Site-Directed Mutagenesis Kit (Thermo Scientific, Rockford, IL) with primers listed in SI Table S6. Prior to PCR amplification, the primers were treated with T4 Polynucleotide Kinase (Thermo Scientific, Rockford, IL) for 5’ phosphorylation according to the manufacturer’s instructions. The site-directed mutagenesis PCR amplification mixture consisted of 23.5 μL of water, 10 μL of 5× Phusion HF buffer, 1 μL of dNTPs (10 mM), 5 μL of each T4 Polynucleotide Kinase-treated primer (5 μM), 0.5 μL of Phusion hot start DNA polymerase (2 U/μL), and 5 μL of HA plasmid of A/chicken/Wuxi/0405005/2013(H7N9) (Ck/WX13) (1 ng/μL). The PCR parameters used for site-directed mutagenesis were as follows: one cycle at 98°C for 30 seconds, followed by 24 cycles at 98°C for 10 seconds, 69°C for 30 seconds, and 72°C for 2 minutes, followed by a final extension step at 72°C for 5 minutes. The PCR products were digested with 1 μl of FastDigest DpnI at 37°C for 15 minutes. The ligation reaction was performed at room temperature for 5 minutes after digestion. The ligation mixture contained 2 μl of PCR products, 2 μl of 5× rapid ligation buffer, 0.5 μl of T4 DNA ligase, and 5.5μl of water. The ligation products (5 μl) were transformed into DH10B Competent Cells (Thermo Scientific, Rockford, IL) following the manufacturer’s protocol. The plasmids were extracted and then used for virus rescue.

### Virus purification

To prepare for biolayer interferometry and glycan microarray analyses, the viruses were purified using sucrose gradient ultracentrifugation. Briefly, allantoic fluids collected from eggs infected with a testing virus were first centrifuged by 4,000 × g for 30 minutes to remove any cell debris. Any remaining cellular debris was then removed further by ultracentrifugation at 4 °C for 30 minutes at 18,000 × g. The virions were subsequently pelleted by ultracentrifugation at 4 °C for 90 minutes at 112,000 × g. The virus was then collected (a ‘milky’ band at around 40% sucrose) and pelleted again after sucrose gradient ultracentrifugation with four layers (30%, 40%, 50%, and 60%). The purified virus was then stored at -80°C until it was ready for use.

### Glycan microarray analyses

The 75 N-linked glycans (52) were printed on slides derivatized with N-hydroxysuccinimide (NHS), as described elsewhere (53). Each glycan was printed in six replicates in a subarray at a concentration of 100 pM in phosphate buffer (100 mM sodium phosphate buffer, pH 8.5). Prior to the assay, the slides were rehydrated in TSMW buffer (20 mM Tris-HCl, 150 mM NaCl, 2 mM CaCl_2_, 2 mM MgCl_2_, 0.05% Tween, pH 7.4) for 5 minutes. A 15 μl aliquot of 1.0 M sodium bicarbonate (pH 9.0) was added to 150 μl of purified virus, and the virus was incubated with 25 μg of Alexa Fluor 488 NHS Ester (succinimidyl ester; Invitrogen) for 1 hour at 25°C. After overnight dialysis to remove excess Alexa 488, the virus HA titer was checked, and the virus was bound to the glycan array. The labeled viruses were then incubated on the slide at 4°C for 1 hour, washed, and briefly centrifuged before being scanned with an InnoScan 1100 AL fluorescence imager (Innopsys, Carbonne, France). Mean relative fluorescent units (RFU) and standard deviation were calculated for six replicates per virus. A threshold of 500 RFU was set to determine background signal.

### Biolayer interferometry assay and data analyses

The virus receptor binding avidities were determined by a biolayer interferometry assay with an Octet RED instrument (Pall ForteBio, Menlo Park, CA). Two biotinylated glycan analogs (3’SLN: Neu5Acα2-3Galβ1-4GlcNAcβ and 6’SLN: Neu5Acα2-6Galβ1- 4GlcNAcβ) were purchased (GlycoTech, Gaithersburg, MD). Three biotinylated glycans Neu5Acα2- 3Galβ1-4(Fucα1-3)GlcNAcβ (sLeX), Neu5Gcα2-3Galβ1-4GlcNAcβ (3’GLN), and Neu5Gcα2-3Galβ1- 4(Fucα1-3)GlcNAcβ (GLeX) were synthesized as previously described using GlcNAcβ-Biotin as starting substrate (52). The glycans were preloaded onto streptavidin-coated biosensors at up to 0.3 µg/ml for 5 minutes in 1 × kinetic buffer (Pall FortéBio, Menlo Park, CA). Each test virus was diluted to a final concentration of 100 pM with 1 × kinetic buffer containing 10 µM oseltamivir carboxylate (American Radiolabeled Chemicals, St. Louis, MO) and zanamivir (Sigma-Aldrich, St. Louis, MO) to prevent cleavage of the receptor analogs by NA proteins of the influenza virus. The association was measured for 30 minutes at 25°C, as described elsewhere (54). To evaluate the binding ability of a virus, we used one high concentration of glycans (0.5µM) with 100 pM viruses to record the endpoint binding response of 30 minutes at 25°C. The threshold for determining positivity in glycan binding was set using the binding response from the negative control, which did not load virus but phosphate-buffered saline (PBS) only. To quantify virus binding avidity, glycan concentrations ranging from 0.05 to 0.5 ug/mL were used. The obtained binding responses were normalized by dividing them by the highest response value obtained during the experiment. Binding curves were fitted using the binding-saturation method, which was implemented in GraphPad Prism 8 software (https://www.graphpad.com/scientific-software/prism/). Normalized response curves were used to calculate the fractional saturation (*f*) of the sensor surface, as described elsewhere (55). The 50% relative sugar loading concentration (RSL_0.5_) is a measure used to quantify the binding avidity between a virus and a glycan. It is calculated at half the fractional saturation (*f* = 0.5) of the virus against glycan analogs. RSL_0.5_ ranges between 0 and 1, and the lower the RSL_0.5_, the stronger the binding affinity between the virus and the glycan analog. Conversely, the higher the RSL_0.5_, the weaker the binding affinity between the virus and the glycan analog.

### Structural modeling

In this study, the HA protein structure of A/Anhui/1/2013(H7N9) (PDB ID 4BSE) with the receptor α2,6-SLN bound to its RBS was employed as the reference template. It’s noteworthy that the HA protein sequence of A/Anhui/1/2013(H7N9) is 100% identical to that of the Ck/WX13 used in our study. For our analysis, the receptor was manually adjusted to produce an H7:Neu5Ac complex. To simulate the effect of V179 versus I179 on Neu5Ac binding, the valine at position 179 was converted to isoleucine using Coot (56). Following the removal of all other small molecules and glycans in the original dataset, both the wild-type and the V179I mutant underwent energy minimization with Phenix (utilizing Phenix.elbow and Phenix.geometry_minimization) (57). Refining both the wild-type and mutant structures was pursued to eliminate any potential biases associated with the algorithm. For a comparative analysis, the two refined structures were aligned using Pymol (via the Pymol.align function) (The PyMOL Molecular Graphics System, Version 1.3, Schrödinger, LLC).

To model Neu5Gc binding to both wild-type and V179I mutant from Ck/WX13, the Neu5Gc moiety from another previously solved H7 HA structure (PDB ID 7TIV, A/equine/NY/49/73 (H7N7)) was superimposed into the above-mentioned H7:Neu5Ac structure using Pymol. The resulting H7:Neu5Gc complex was subjected to V179I mutation, energy minimization and comparative analysis using the same modeling procedures mentioned above.

All structural figures were prepared using Pymol (The PyMOL Molecular Graphics System, Version 1.3, Schrödinger, LLC).

### Growth kinetics in Neu5Gc expressed MDCK cells

To evaluate the impact of Neu5Gc expression on virus infectivity of H7 viruses, we conducted growth kinetics analyses of three selected viruses (rgMall/NJ10, rgCk/WX13, and rgCk/HN17) on both MDCK-wt and MDCK-Gc. Additionally, we included two mutant viruses (A122N and V179I) in the growth kinetics analyses to assess whether each of these three amino acid substitutions may affect virus replication efficiency. To initiate infection, cells were seeded in 6-well plates and allowed to grow for approximately 18 hours, reaching 90% confluency. The cells were then infected with each testing virus at a multiplicity of infection (MOI) of 0.001. Following infection, supernatants were collected at 12, 24, 36, and 48 hours post-infection and subjected to viral titration.

### Viral titration

For viral titration, we determined the 50% tissue culture infection dose (TCID50) on MDCK CCL-34 cells. Briefly, cells were seeded at a density of 2 × 10^4^ cells per well in a 96-well plate with Opti-MEM I Reduced Serum Medium. Cells were then incubated at 37°C with 5% CO_2_ for 18-20 hours before virus inoculation. Viral samples were serially diluted in Opti-MEM I Reduced Serum Medium supplemented with 1 µg/mL of TPCK-trypsin. Subsequently, 200 µL of each virus dilution was inoculated onto MDCK cells in quadruplicate. Infected cells were then incubated at 37°C with 5% CO_2_ for 72 hours and evaluated for positivity using hemagglutination assays. The number of positive and negative wells for each dilution were recorded for TCID50 calculation based on the method described by Reed and Muench (58).

### Hemagglutination assays

Hemagglutination assays were carried out by using 0.5% turkey erythrocytes as described elsewhere (59).

### Characterization of adaptative mutations on cells

To investigate the effects of Neu5Ac and Neu5Gc on the adaptive mutations, we passaged rgCk/WX13 and rgMall/NJ10 in MDCK-wt and MDCK-Gc cells five times. For each passage, the original seed virus or supernatants were diluted 200-fold during the virus infection. Viral RNA was extracted from the seed virus and supernatants from the fifth passage and subjected to next-generation sequencing.

### Next generation sequencing, genomic assembly, and polymorphism analyses

Conventional two-step RT-PCR whole genome amplification was set up using 8 pairs of universal primers (50). Viral RNA was reverse transcribed to cDNA using SuperScript III Reverse Transcriptase (ThermoFisher Scientific, Cat. #: 18080-044). PCR amplification was subsequently performed using Platinum Tag DNA Polymerase High Fidelity (ThermoFisher Scientific, Cat. #: 11304-102). Amplicons of the same samples were pooled together before purification. AMPure XP beads (Beckman Coulter, Cat. #: A63881) purified amplicons were analyzed for cDNA quality and quantity using TapeStation 4200 (Agilent Technologies, Santa Clara, CA) DNA5000 kit (Agilent Technologies, Cat. #: 5067-5588, 5589).

After TapeStation analysis, ∼50 ng of pooled cDNA of each sample was used as input for library preparation using illumina DNA prep kit (illumina, Cat. # 20018705) following the manufacturers’ instructions. The purified libraries with different indexes were quantitated using TapeStation D5000 kit and equal molar of each library were pooled together. Pooled libraries were denatured, diluted to an appropriate loading concentration and loaded onto Miseq 600 cycles V3 cartridge (illumina, Cat. #: MS-102-3003) for sequencing.

Iterative Refinement Meta-Assembler (IRMA) v.1.0.3 (https://wonder.cdc.gov/amd/flu/irma/) was used for sequence assembly and nucleotide variant analysis, and the results were further validated by CLC Genomics Workbench v21.0.3. The quality of the reads was trimmed with a Phred quality score of 20, which indicates a base call accuracy of 99%, the likelihood of finding one incorrect base call among 100 bases. The polymorphisms were analyzed by using DiversiTools (http://josephhughes.github.io/DiversiTools/). The most abundant nonsynonymous mutations in HA protein were plotted to visualize adaptive amino acid substitutions caused by cell passages.

To identify intra-host genetic diversity of H7 viruses, minor amino acid variants ratio was calculated by the sum of the second and third amino acid counts divided by the sum of the top three amino acid counts in each residue. Ggplot R package was used for amino acid variants visualization. To minimize the impacts of potential sequence errors on the intra-host variant analyses, only those amino acid variants ratio with >5% were considered.

### Multiple sequence alignment and phylogenetic analyses

Multiple sequence alignments were generated using Muscle v5.1 (60). The approximately-maximum-likelihood phylogenetic tree was inferred by using fastTree v 2.1.11 (61). Phylogenetic trees were visualized by using FigTree v1.4.3 (http://tree.bio.ed.ac.uk/software/figtree/).

### Characterization of Neu5Gc expression in avian respiratory and gastrointestinal tracts

A total of seven species were studied in this study, including chicken (*Gallus gallus*), Canada goose (*Branta canadensis*), mallard (*Anas platyrhynchos*), gadwall (*Mareca strepera*), green-winged teal (*Anas carolinensis*), northern shoveler (*Spatula clypeata*), and wood duck (*Aix sponsa*). The avian respiratory and gastrointestinal tract (cloaca and colon) tissues were collected and fixed by submerging them in 10% neutral buffered formalin, and then they were embedded in paraffin. Sections of 5 μm were made from the embedded tissues. The tissue sections were deparaffinized by dipping into the following solutions: 3 times of 10 minutes in xylene, 3 minutes of 100% ethanol, 100% ethanol, 95% ethanol, 70% ethanol, 50% ethanol and rinsed in ddH_2_O. Antigen was then heat-induced target retrieved with diluted Target Retrieval Solution, Citrate pH 6.1 (10x) following the manufacturer’s manual (Dako, Carpinteria, CA). The sections were blocked by 3% Bovine Serum Albumin for 1 hour at room temperature. The sections were rinsed with PBS and incubated with the anti-Neu5Gc (1:500 dilution; Biolegend, San Diego, CA) overnight at 4°C. Sections were incubated with goat anti-chicken IgY (H+L) secondary antibody conjugated with Alexa Fluor™ 594 at 1:500 dilution with PBS before counterstaining with DAPI. The sections were washed three times of 5 minutes with PBST after every step of antibody incubation. The slides were air dried and covered with coverslips by using Prolong antifade reagent. Images were captured with the Zeiss Axiovert 200M.

### Structural visualization

The HA protein was visualized in PyMOL using the template HA protein structure of A/Shanghai/02/2013(H7N9) (accession number: 4LN3) from the Protein Data Bank (PDB, https://www.rcsb.org/).

### Data availability

We have submitted the raw and assembled genomic data collected from this study to GenBank with the BioProject accession number PRJNA978106. This submission includes six datasets (seed virus rgCk/WX13 and Mall/NJ10 and their corresponding 5th passages in MDCK-wt and in MDCK- Gc), datasets for 85 H7 viruses from dabbling ducks, and 18 H7 viruses from domestic poultry from North American.

### Statistical analyses

A two-way ANOVA test was performed using GraphPad Prism 8 (https://www.graphpad.com/scientific-software/prism/) to compare the statistical differences between viral titers at different time points in the growth kinetics of both the wild type and a testing mutant. A P-value of 0.05 was considered significant.

## Acknowledgments

We are thankful to Wendy S. Weichert for technical support and appreciate comments on prior drafts of this manuscript provided by Andrew Ramey. We would like to acknowledge the useful resource of Influenza Research Database and the support from BEI resources. This project was partially supported by the National Institutes of Health (grant number R21AI144433), the National Science Foundation (#2109745), and the Welch Foundation (C-1565 to YJT).

The findings and conclusions in this report are those of the authors and do not necessarily represent the official position of the U.S. Government.

## Supporting information captions

### List of Supplementary Tables

**Table S1.** Distribution of H7 influenza A viruses (IAVs) in avian species.

**Table S2.** Amino acid polymorphisms among H7 viruses from public databases.

**Table S3.** List of H7 avian influenza viruses used in the intra-host genomic diversity analyses.

**Table S4.** Intra-host amino acid polymorphisms for H7 avian influenza viruses

**Table S5.** List of glycans printed on the glycan microarray.

**Table S6.** List of forward (F) and reverse (R) primers used in generating HA mutants by site-directed mutagenesis.

### List of Supplementary Figures

**Fig. S1.** Distribution of H7 influenza A viruses (IAVs) in avian species.

**Fig. S2.** Endpoint binding analyses of virus-glycan interaction using biolayer interferometry for H7 mutant viruses.

**Fig. S3.** The H7 IAVs detected recently in the US domestic poultry gained limited adaptation to the poultry host.

## Notes

### Competing Interest Statement

The authors have declared no competing interest.

